# MCO: towards an ontology and unified vocabulary for a framework-based annotation of microbial growth conditions

**DOI:** 10.1101/218289

**Authors:** VH Tierrafría, C Mejía-Almonte, JM Camacho-Zaragoza, H Salgado, K Alquicira, S Gama-Castro, J. Collado-Vides

## Abstract

**Motivation:** A major component in our understanding of the biology of an organism is the mapping of its genotypic potential into the repertoire of its phenotypic expression profiles. This genotypic to phenotypic mapping is executed by the machinery of gene regulation that turns genes on and off, which in microorganisms is essentially studied by changes in growth conditions and genetic modifications. Although many efforts have been made to systematize the annotation of experimental conditions in microbiology, the available annotation is not based on a consistent and controlled vocabulary for the unambiguous description of growth conditions, making difficult the identification of biologically meaningful comparisons of knowledge generated in different experiments or laboratories, a task urgently needed given the massive amounts of data generated by high throughput (HT) technologies.

**Results:** We curated terms related to experimental conditions that affect gene expression in *E. coli* K-12. Since this is the best studied microorganism, the collected terms are the seed for the first version of the Microbial Conditions Ontology (MCO), a controlled and structured vocabulary that can be expanded to annotate microbial conditions in general. Moreover, we developed an annotation framework using the MCO terms to describe experimental conditions, providing the foundation to identify regulatory networks that operate under a particular condition. MCO supports comparisons of HT-derived data from different repositories. In this sense, we started to map common RegulonDB terms and Colombos bacterial expression compendia terms to MCO.

**Availability and Implementation:** As far as we know, MCO is the first ontology for growth conditions of any bacterial organism and it is available at http://regulondb.ccg.unam.mx/. Furthermore, we will disseminate MCO throughout the Open Biomedical Ontology (OBO) Foundry in order to set a standard for the annotation of gene expression data derived from conventional as well as HT experiments in *E. coli* and other microbial organisms. This will enable the comparison of data from diverse data sources.

## Introduction

As any other living organism, *E. coli* needs to be able to keep a constant monitoring of their surroundings in order to rapidly respond and adapt its physiology to an ever-changing environment and be able to thrive and survive. To achieve this, *E. coli* has developed a set of mechanisms and signaling pathways to sense different kinds of stimuli, and to transduce these external cues into the appropriate physiological response by adjusting the patterns of gene expression. In this way, transcriptional regulation and growth conditions are fundamentally related. Additionally, *E.coli* has been long considered a model organism mainly because of the vast amount of biological information that has been gathered through decades. Much of this information is now available through high-quality manually-curated knowledge bases, which enable a systems-level analysis, modeling, and mechanistic interpretation of the phenotypic behavior by computational approaches.

The integration of growth conditions in genome-scale computational models of transcriptional regulation and metabolism is essential, since even a single change in one of the factors of the growth conditions can result in huge changes in the patterns of gene expression. In this sense, MediaDB [1] recognizes the importance of growth conditions providing an invaluable compendium of media chemical composition, in addition to other important physiological information such as growth rates and secretion/uptake rates for several organisms under different conditions. On the other hand, RegulonDB currently gathers the most updated and comprehensive collection of mechanistic knowledge of transcriptional regulation in *E. coli*, curated from original scientific publications [2]. Since 2013, RegulonDB started collecting information related to growth conditions, further enriching the content and significance of the provided data [3]. Moreover, repositories such as GEO in compliance with MIAME guidelines [4], have adopted Minimum Information Standards for the submission procedures of datasets of high-throughput experiments, which include the annotation of the experimental factors of the different samples. However, one of the major challenges to fully harness the potential of knowledge integration is the consistency of the vocabulary used to describe growth conditions.

In an effort to standardize the annotation of experimental conditions in microbial data repositories, here we describe the development of the Microbial Conditions Ontology (MCO). MCO is a domain ontology which provides a controlled vocabulary to describe microbial growth conditions, built on top of the standard upper level Basic Formal Ontology (BFO) [5]. This ontology provides growth conditions terms together with their definitions, synonyms, references, and higher-level relations, that can unambiguously define and tag each attribute of a particular experimental condition in order to systematize the annotation. Furthermore, the implementation of this ontology in RegulonDB represents another step towards an efficient, accurate and inter-operable retrieval, comparison and analysis of biological information in accordance with the recent call to follow the principles of the FAIRification process of data (Findable, Accessible, Interoperable, Reusable) [6, 7]

Currently, there are two ontologies related to prokaryotic organisms: the Ontology for Microbial Phenotypes (MPO) [8], and the Ontology of Phenotypic and Metabolic Characters (MicrO) [9]. While MPO aims at language standardization to describe bacterial phenotypes, MicrO aims at capturing evolutionary diversity and at using logical inference to automatically populate some character matrices. Since a fundamental aspect of bacterial diversity lies in the metabolic chemical transformations, MicrO makes formal definitions that relate phenotypic traits with growth media composition and metabolic processes. On the other hand, Growth Medium Ontology (GMO) was developed as a controlled vocabulary to describe ingredients that constitute microbial growth media and to annotate Japanese biological bioinformatic resources, but so far it does not include composition nor definitions [http://bioportal.bioontology.org/ontologies/GMO]. Ontologies related to growth conditions are: the Exposure Ontology (ExO), aiming to link environmental contributions to human diseases [10]; the Microarray Experimental Conditions (MO), aiming to standardize the description of microarray experiments (http://mged.sourceforge.net/ontologies/index.php); the Experimental Conditions Ontology (XCO), aiming to describe conditions under which physiological and morphological measurements are made in studies involving humans or model organisms [11]; Plant Environment Ontology (EO), aiming to describe growth conditions, types of experiments and treatments in plant biology experiments [12]; and the Zebrafish Conditions Ontology (ZECO), aiming to describe the zebrafish experiments conditions (https://github.com/ybradford/zebrafish-experimental-conditions-ontology). Hence, although some components to describe bacterial growth conditions are included in other ontologies, there is not yet an ontology adequately to address growth conditions and strains used in microbial experiments studying gene regulation.

## Methods

### Gathering terms that describe growth conditions

The initial set of terms describing growth conditions was obtained from RegulonDB version 9.4 [2] and Colombos version 3.0 [13]. From RegulonDB we considered three datasets: former GCs, effectors of transcription factors (TFs), and TFs summaries. The first dataset contains the experimental variables amongst the control and experimental test. On the other hand, the effectors dataset includes some compounds that affect the active/inactive conformation of cognate TFs, while the TFs summaries contain GCs data related to the expression of genes encoding TFs or those related to the activation or inactivation of TFs function. The terms obtained from Colombos depict the GCs under which high throughput (HT) experiments, including microarray and RNA-Seq, of several prokaryotic species were done.

Additional terms were obtained from 43 papers according to the items previously proposed by Frederick Neidhardt (strain, medium, aeration, temperature, growth phase and growth rate) [14], and by our research group (medium supplements, pH, pressure and optical density -OD-). Taking together these elements, we further developed a framework that specifies the minimal information required to describe growth conditions, while satisfying a description that guarantees reproducibility. However, to achieve this, it was necessary to make a slight modification consisting in replacing the *strain* component name, by *genetic background*, because the strain name does not bring precise information about genetic modifications such as knock-out of genes, which are frequently used in this type of experiments.

Considering the aspects mentioned above, the resulting framework was composed by the following items: 1) genetic background, 2) medium, 3) medium supplements, 4) aeration, 5) temperature, 6) pressure, 7) pH, 8) OD, 9) growth phase, and 10) growth rate. Written in this order, we assume that this framework will describe the growth conditions evenly and consistently, and will make easier the identification of those experiments performed under similar conditions for the sake of relevant biological knowledge.

### Ontology development

As described above, the description of growth conditions involves different pieces of information. Thus, there are two possibilities to create terms to annotate growth conditions. The first one is to create fully composed ontological terms; these terms would be phrases including all of the required pieces of information. Annotation with composed terms is direct, as one term fully describe a growth condition. The second one is to create simple ontological terms to describe elementary components. These terms are combined at the time of annotations. In other words, annotations of one growth condition requires one or more simple terms to be fully described. We used this last approach called post-composition [15].

We proceeded in a bottom-up approach in the ontology development by gathering specific terms used to describe growth conditions, followed by a classification and hierarchy construction stage [16]. Classification included synonym sets definition. To ensure interoperability with other resources, our ontology was developed under the upper-level Basic Formal Ontology (BFO) [5].

Here we refer as concepts to defined entities that may have alternative names or synonyms. Having sketched the hierarchy of required concepts, we proceeded to search for terms to refer to these concepts in extant ontologies. Aiming to be part of the OBO Foundry, we did this search using OntoBee [17]. In some cases, more than one keyword was used to search in these ontologies for a given concept, since each concept might have one or more synonyms. The result of this search was a set of ontologies and a set of Internationalized Resource Identifiers (IRIs) of the required concepts.

To extract and merge the required classes from these ontologies into MCO, we used the ontology management command line tool ROBOT [18]. For chemical terms, we extracted their whole ChEBI classification [19]. For other terms, we used MIREOT method [20] to extract shorter hierarchies or specific terms that would allow us to build our own hierarchy using little pieces of diverse ontologies.

Some chemical terms of our initial set were not included in ChEBI, thus we requested their incorporation. For all other kinds of growth conditions terms that were not found in other ontologies, we created our own terms. This was because, to our knowledge, ChEBI is a well consolidated ontology that has a precise scope in the sense of the kinds of entities it represents, but it is completely organism-independent. In contrast, the other reused ontologies are less well known and have broader scopes in the kind of entities they represent, but narrower scope since they may be organism-specific (see Results for specific reused ontologies).

Finally, MCO version 1.0 was created using Protégé version 5.1 [21, 22] in owl format, based on OBO principles [23]. We are currently discussing our ontological model with members of the OBO foundry in order to incorporate MCO to their set of standards. To programmatically analyze and edit the ontology, we used the python library owlready [24].

## Results

### Manual curation of growth condition terms according to an annotation framework

Initially, a collection of 424 terms was recovered from former GCs, effectors and TF notes in RegulonDB. These terms are predominantly defined by the experimental variable, that is to say, the metabolite or physical condition that is added in the experiment and is absent in the control. However, GCs can be described by further information as stated in the Methods section, i.e. genetic background, medium, medium supplements, pH, aeration, temperature, pressure, OD, growth phase and growth rate. Thus, we realized that we were not recovering all the available information regarding growth conditions. Consequently, these elements were considered to better describe the experimental conditions used in laboratory. We manually curated a set of 43 papers originally associated with terms where the experimental treatment was not known or ambiguous, such as "with antibiotic stress", "growth on non-optimal carbon source" and "growth with metal".

In the following, we report details of curation for each element of the proposed framework, providing examples of how we built the current controlled vocabulary. This will offer a summary of the type of decisions made, sometimes as a balance between the theoretical ideal in face of the facts of how this information is reported. This will also provide a sample of the quality and precision used in reporting the experimental work. Our curation involved describing growth conditions terms for 598 total experiments, including both control and experimental tests. We also report the resulting classification of the recovered words into the ontology, beginning by stating under which class of BFO the terms fall into.

### Genetic background

Regarding the genetic background component, different authors use different ways to refer to gene deletion, including simple ways such as “gene-”, “∆gene”, “gene mutant”, and more complex ways to indicate genetic modifications such as those that indicate the name of an antibiotic resistance gene or a transposable element (Tn) used to replace a particular gene, for instance: “gene::kan” and “gene::Tn1”, respectively. Moreover, some authors simply put a name to their mutant strains “JA173”. This shows that there is a great number and diversity of terms to refer to a deletion mutant. Since the aim of all these examples is the inactivation of gene activity, we unified this information under the "knockout mutant" term, thus 45 terms were obtained.

In order to indicate a point mutation, we propose firstly to identify the type of mutation, i.e. insertion, deletion or substitution, and secondly to annotate it in relation to the translational start site of the corresponding gene. In this way, we built the following term “C-T transition at nt −10 from bioA translational start site”. This was done to make such piece of information depend only on the protein information because published papers describe point mutations typically referring to a small fragment of DNA that carry its own numbering, usually different from that of the genome sequence.

Compared to gene deletion or point mutations, mutant complementation is much more complex, given the number of strategies that can be used to restore gene deletion, which involve the use of replicative- (either single- or multicopy), integrative- or inducible plasmids (including their inducers), even when the inducer is not added to the culture to prevent the expression of bearing gene(s), a technique commonly used in control assays. In this scenario, and to evenly describe all these genetic variants, it was necessary to establish a syntax that considers the following parameters when plasmids are used: the copy number of the plasmid, the *plasmid* word, the vector name, the plasmid inducer followed by the *induced* word, and the cloned gene or genes. If plasmids were not induced, it is not necessary to mention the inducer name, but the term must include the *not-induced* word (Table 1). This notation has also the advantage of enabling heterologous mutations to be properly described, for instance: "plasmid pSKOG (*K. oxytoca hydG*)". Using this syntax, we obtained 33 terms.

**Table 1.**
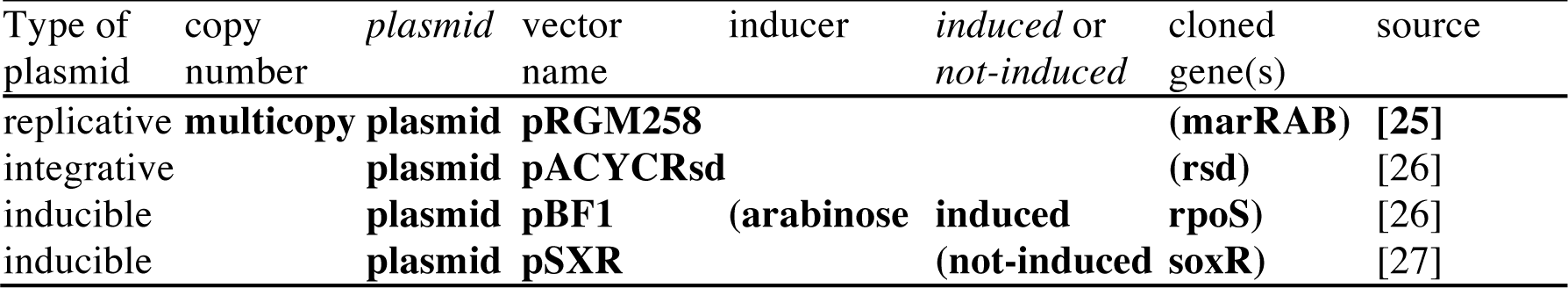
Selected terms constructed according to the proposed syntax to describe plasmid-mediated genetic complementation

In addition to plasmids, phages and transposons are also used to restore gene deletions, however, these genetic tools are less frequently used. According to this, we found only four terms that involve genetic complementation mediated by either phages: "Φ(soxR)", "λ(soxR)" and "λ(truncated soxR)"; or transposons: “mini-Tn5-nrdEF transposon”.

Last but not least is the “wild type” term, thus, considering all terms used to describe gene deletions, point mutations, mutant complementation, as well as the wild type background, we collected a total of 84 terms associated with genetic background (Table 2).

**Table 2.**
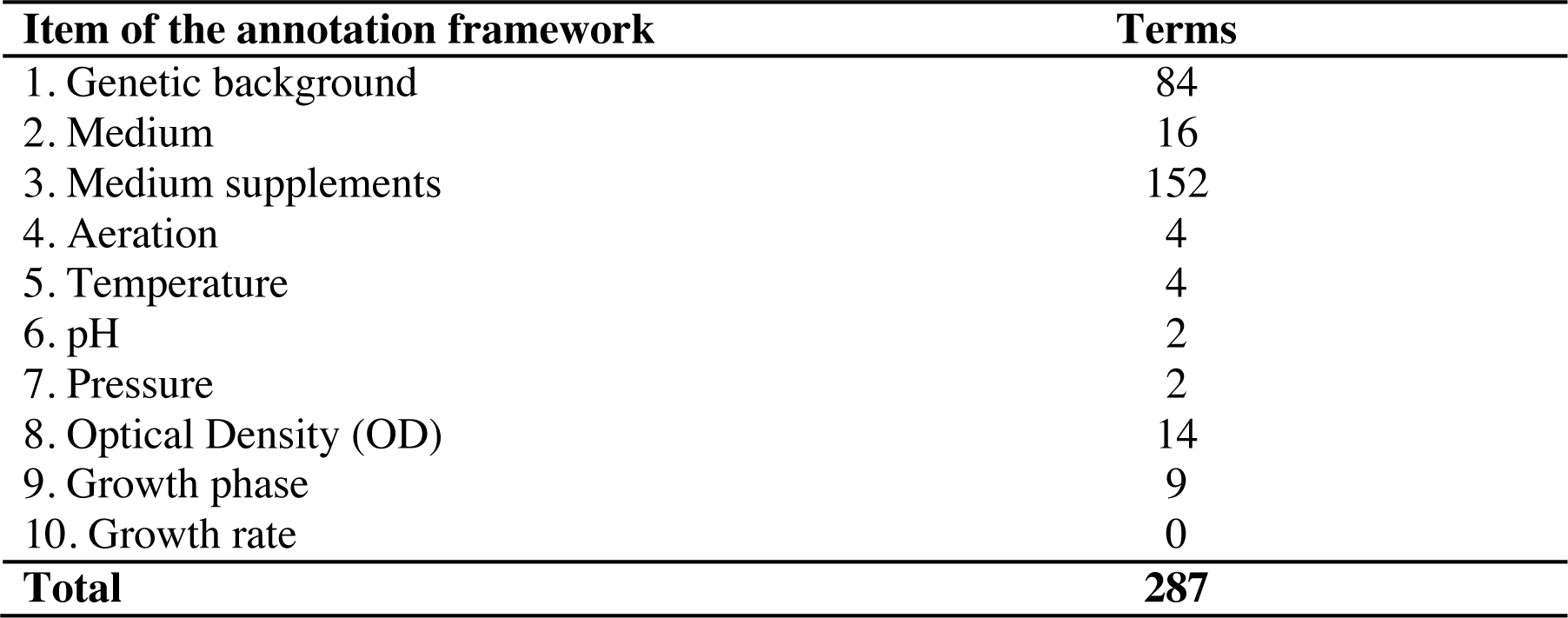
Growth conditions-related unique terms obtained from manual curation of 43

Genetic background is described with BFO qualities. A quality in the framework of the Basic Formal Ontology is a dependent continuant, which means that qualities only exist if the bearer of the quality exists. In words of the BFO’s authors, dependent entities inhere in substantial entities, inherence being an existential dependence [5].

A quick review of the genotype concept in OBO ontologies shows that it has been regarded as an object in SO [28], SIO [29] and ExO [11]; an object aggregate in OGI; a quality in OMIT [30]; and a generically dependent continuant (entities whose realization depends on material entities, but exist independent of time and space) in GENO (Lin 2009; Brush 2013). Because we will not annotate sequence features, we consider that the best way to define it is as a biological quality inherent to the bearer (genome, cell, organism) by virtue that any life form has genetic content. A genotype is a quality of genomes that describes genetic variation. This allows us to describe cell cultures used in the experiments in terms of two kinds of genotypes: mutant and wild type. We found terms describing types of mutations in OntoBee, but not the terms to describe mutants as genotypic qualities. Thus, we created a hierarchy of mutant genotypes.

We have two classification schemes of mutants: one based on a structural criteria, and the other based on the effect of the mutations. Structurally defined mutant genotypes are represented in four classes which may not be disjoint: episomal expression mutant, gene variant mutant, non-coding region variant mutant and insertion mutant. Mutant genotypes defined by the their effects are represented in three classes which are disjoint: knockdown mutant, knockout mutant and overexpression mutant (Fig. 1). The most specific classes describe gene-specific mutants.

**Figure 1.**
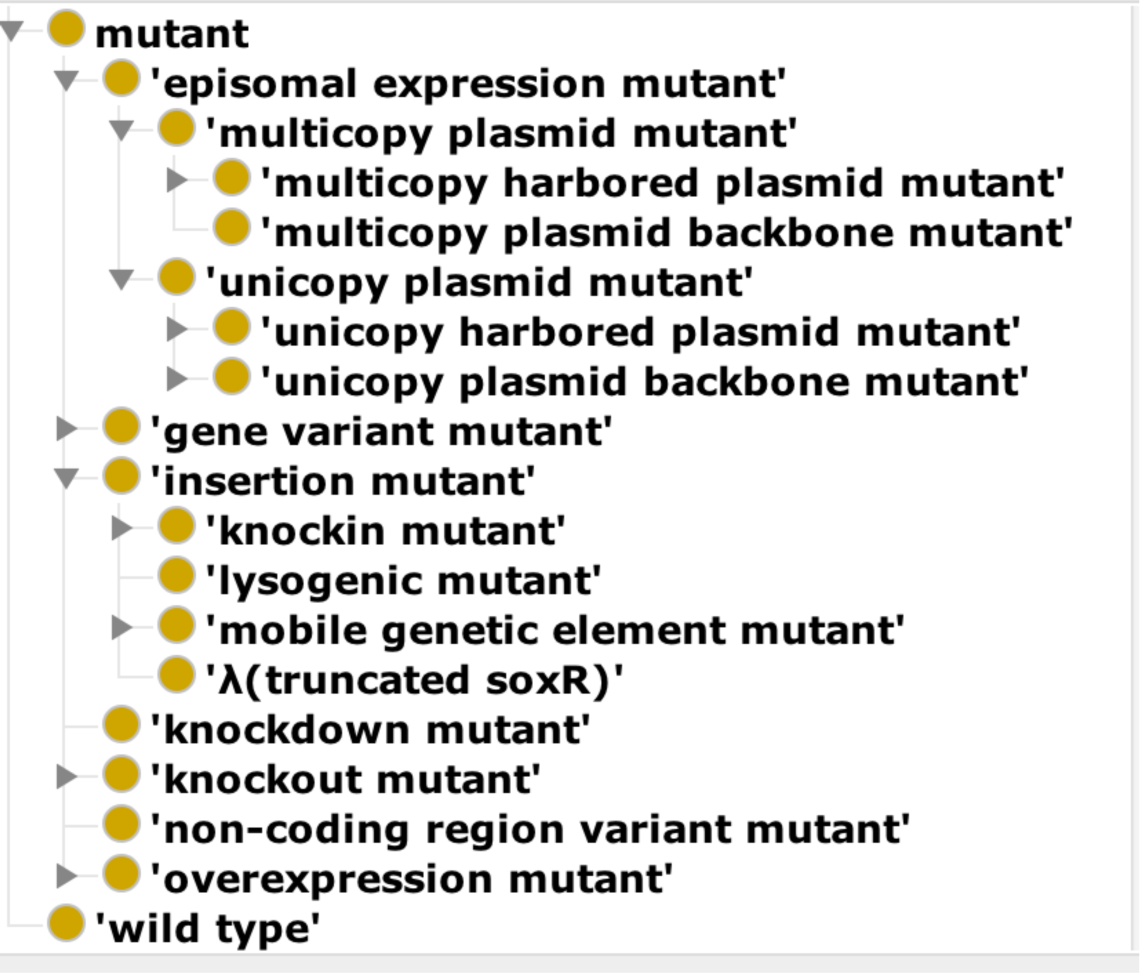
Mutants classification according to MCO

If we base mutant annotation on structural criteria, there will be multiple terms referring to one single effect, since one mutation effect can be achieved by different methods. For example, a knockout mutant can be created by deletion of the gene of study or interruption by insertion of the coding sequence of the gene. Thus, our annotation framework mandatorily requires a term describing the effect and, optionally, a term describing the structural nature of the genetic modification. Annotation of the effect is fundamental since this is what ultimately determines the physiological outcome of the mutation. On the other hand, annotation of structural details is important specially in the case of episomal expression, since it is known that copy number variations of plasmids can result in a different metabolic burden, which may have phenotypic consequences.

### Culture medium

We found 16 terms related to the medium object (Table 2) including “LB” and several variants of minimal media, i.e. “MM”, “M9”, “MOPS”, and “M56”, as expected. Other media used to cultivate *E. coli* include: “MacConkey lactose agar”, casein acid hydrolysate “CAA”, “TGYEP”, and a medium devoid of nitrogen and carbon sources, the “N^-^ C^-^ medium”. Additionally, we found different conditioned media, that is to say, the supernatant from a culture grown until a specific OD, derived either from a wild type: “conditioned medium”, or from a particular strain: “LE392 conditioned medium”. In fact, we found that some authors added a particular solvent to a basal or conditioned medium to extract and probe the effect of extracted components, accordingly, we discovered the terms “ethyl acetate extract of LB” and “ethyl acetate extract of conditioned medium”. In contrast to the complexity to describe these media, we encountered simpler terms such as “rich medium” and “poor medium”.

The “MOPS” and “MacConkey lactose agar” terms deserves special attention. Although they strictly refer to the addition of 3-(N-morpholino) propanesulfonic acid, or lactose to the medium to maintain stable near-neutral pH-levels, or select *E. coli* while growing, respectively, and thus these compounds may be considered as medium supplements (see next section); the scientific literature regularly refers to those media simply as described above.

Culture media belong to the class object of the BFO. We made a classification of culture media that currently does not exist in other ontologies. The most general classes are natural medium and artificial medium. Natural medium takes classes from UBERON (Uberon Multispecies Anatomy Ontology) [31], CL(Cell Ontology) [32] and CLO (Cell Line Ontology) [33]. These classes refer to body fluids and kinds of cells that are used as culture media in some experiments. Artificial medium refers to culture medium specifically made to grow microorganisms. The next level of classifications is related to the nutrient content: minimal, poor and rich media. We think this classification is more relevant for the purposes of our ontology because the class to which a determined medium belongs to, tells us if the bacteria were subject to a nutrient stress condition or not, which might have a significant impact on the regulatory and phenotypic outcome. This classification will also allow us to make ontological relations with the higher level terms that describe stress conditions, frequently referred to in the literature by microbiologists (see Inferences section).

Under this higher classification of media, we added classes to specific defined media (e.g. LB and M9 minimal medium). Some of these terms were already found in other ontologies such as GMO [http://bioportal.bioontology.org/ontologies/GMO] and MicrO. However, we did not import those terms since not all media we curated were included in those ontologies, they lacked a full description of the media composition and definition, or we could not process the original ontology with ROBOT owing to illegal reuse of entities.

### Medium supplements

“Medium supplements” was the most diverse aspect we studied, reflecting the vast repertoire of treatments used to analyze gene expression. This item comprises 152 terms (Table 2), 15 of which are one-word terms that simply indicate the name of a particular compound added to the medium, for instance “glucose”, whereas 132 terms indicate the usage concentrations, which additionally can be expressed in different units, for example, we found that glucose was added at: 0.05, 0.2, 0.4, 0.5 or 2 %; 10, 11, 60 or 120 mM; and 2 or 4 g·L^−1^. Additionally, a small group of 5 terms were used to annotate a range of concentrations. This term are: “ZnCl_2_ 0.2 to 1 mM”, “Hg(II) 1 to 10 µM”, “Cd(II) 1 to 100 µM”, “trehalose 0.5 to 0.7 M”, and “trehalose above 1.2 M”.

Finding a diversity of terms referring to medium supplements convey some obstacles. For example, not all the collected chemicals were deposited in the ontologies that support MCO, specifically in ChEBI. This was the case of the bile salt “sodium ursodeoxycholate” (NaUDC), cytidine triphosphate “CTP”, “trehalose”, “potassium glutamate”, “*V. harveyi* autoinducer” (N-(3-hydroxybutanoyl)-L-homoserine lactone), N-Decanoyl-DL-homoserine lactone “DHSL”, as well as the “fumarate” and “aspartate” ions. For the majority of these compounds, solely the requesting to ChEBI ontology for their incorporation may be sufficient. Nonetheless, we faced a particular difficulty in the annotation of fumarate or aspartate because ions, as such, cannot be added to the medium, they need to be added to the medium either in their salt or acid form, i.e. fumaric acid or sodium fumarate. For this reason, it is not possible to infer the reagent used if it is not specified by the authors. Moreover, from an ontological perspective, “fumarate” and “aspartate” are general classes that include several ions, i.e., fumarate(−1) and fumarate(−2). Despite this, the sole mention of generic ion names is a common practice, thus we decided to preserve these general terms to refer to diverse ions (fumarate). Ultimately, it is important to note that we annotated the name of commercially available reagents if authors mentioned it at least once, thus the precise salt may be found.

Supplements belong to the class object in the framework of BFO. Since these are chemicals, we used ChEBI terms to describe this growth condition feature. We selected only those chemical compounds that have been used as additives in *E. coli* cultures. Currently there are 2074 ChEBI classes, including not only specific molecules, but also their classification.

### Physical qualities

As mentioned earlier, qualities are entities inherent to substances. In this sense, the following elements: aeration (the amount of oxygen or air the bearer contains), temperature (the amount of thermal energy the bearer has), pressure (the force exerted by the bearer per unit area), pH (the amount of hydrogen ions contained by the bearer) and OD (the amount of light the bearer is able to transmit) are physical qualities inherent to the growth medium.

Each of both aeration and temperature features is composed by four elements: “oxygen”, “oxygen 10 µM”, “oxygen > 10 µM”, and “oxygen < 10 µM”; and “37 ºC”, “30 ºC”, “28 ºC”, and “32 ºC”, respectively. On the other hand, pressure and pH aspects contain only two terms each: “0.1 MPa” and “10 to 40 Mpa”; and “pH 7.0” and “pH 6.5”, respectively (Table 2). Although these two parameters contain the same amount of terms, it is important to note that there is a significant difference in the availability of this information. In this sense, we found pressure-related information only in 4 out of 598 experiments (less than 1 %), whereas, relative to pressure, the pH data was 10-times more reported, with 42 experiments (Fig. 2).

**Figure 2.**
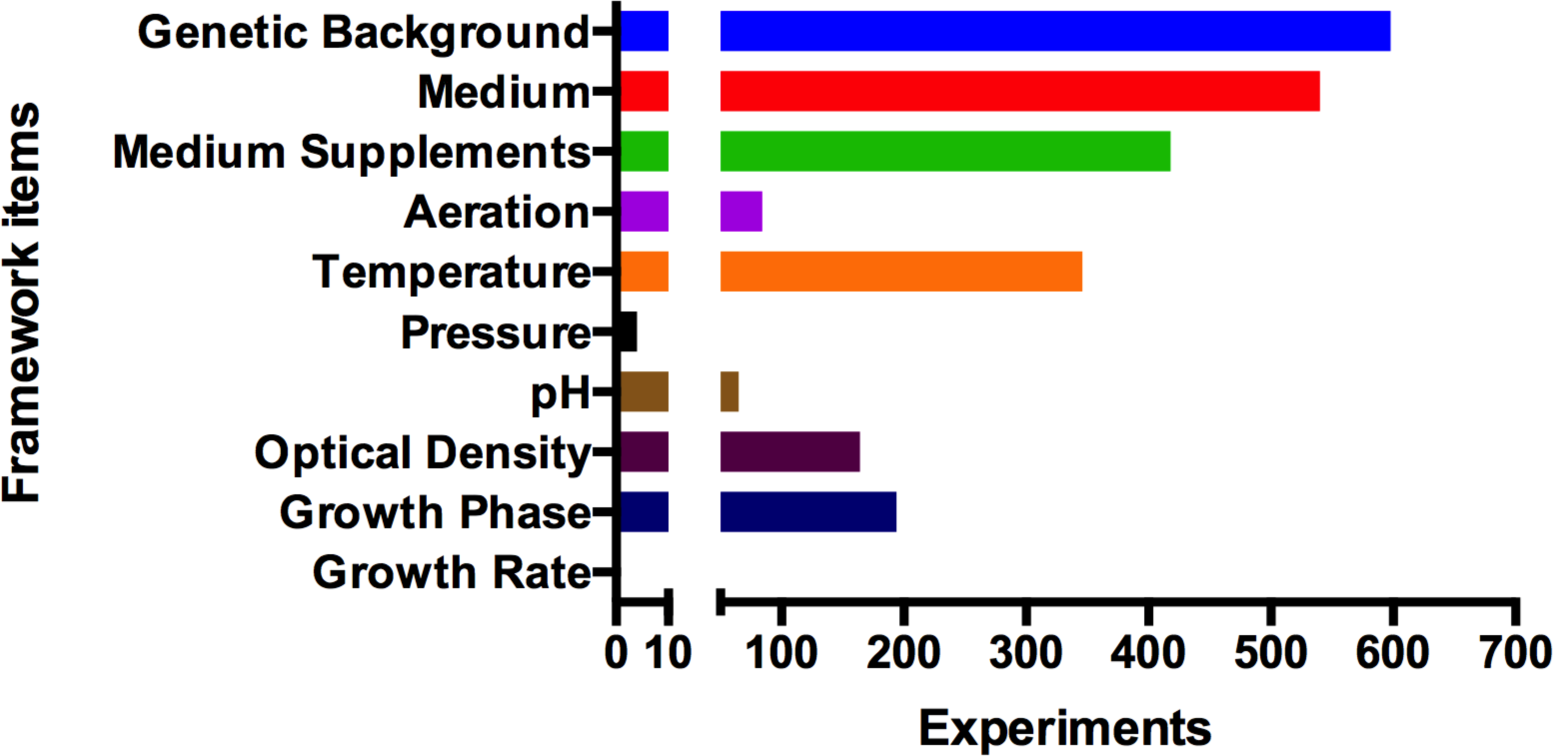
Distribution of information on growth conditions reported in 598 experiments revealed by manual curation performed in this study

In addition to physical qualities just described, we found out 14 terms related to OD (Table 2). Ten of these terms indicate a specific OD value, for instance, “OD600 0.5”, while the remaining 4 terms indicate intervals such as “OD650 between 0.5 and 0.7”. Noteworthy, we detected several terms indicating overlapping intervals, including: “OD600 below 0.7”, “OD600 0.4 to 0.5” and “OD600 0.4 to 0.6”.

The majority of classes to represent these features in the ontology were imported from other ontologies. We added PATO’s physical quality class as a child node of BFO’s quality. Cellular motility, pressure, and temperature were taken from PATO (Phenotypic Quality), atmospheric pressure from EO (Plant Environment Ontology), oxygen content and pH from ZECO (Zebrafish Experimental Conditions Ontology), and gravitation from OMIT (Ontology of MIRNA Target) [15]. We created terms for optical density and aeration, since we did not find them in other ontologies. We found the process of measuring optical density in CHMO (Chemical Methods Ontology) [https://github.com/batchelorc], but not optical density as a quality. We made oxygen content subclass of aeration since aeration was defined as the quality of containing either oxygen or air.

### Growth

This section includes both the growth rate, and growth phase elements. In some theoretical frameworks, growth rate is considered a fundamental property, particularly in rapidly reproducing organisms as bacteria, because cell size and macromolecular composition, which are collectively referred to as the physiological state of the cell, vary with growth rate [34]. In other words, the physiological state is influenced by the growth rate that nutritional conditions support, not only by the specific nutrients in the medium, i.e. cell cultures growing in different media composition that are growing at the same rate have the same physiological state. Even though it makes all sense theoretically, we did not find any term related to this aspect (Fig. 2), revealing that, at least in the 43 papers reviewed, authors rarely report it.

Moreover, the only reproducible state occurs after cells have completed their chemical adjustments to a specific growth condition and the exponential increase in the mass of the population occurs at a constant rate (exponential phase), and before they change the medium as a result of growth. This state is called steady-state because every cellular component increases by the same constant factor per unit time [14].

Otherwise, author frequently specify the growth phase where gene expression was analyzed. We found 194 experiments indicating this parameter, therefore, growth phase was the major-used element to report any aspect related with growth, including OD or growth rate (Fig. 2).

The experiments that reported the growth phase accommodated in 9 terms (Table 2) indicating, similar to the OD terms, either a punctual- or located within an interval growth phase. Accordingly, we collected 6 terms used to indicate a specific growth phase such as “exponential phase”, “mid exponential phase”, “stationary phase”, “late exponential phase”, “early stationary phase” and “early exponential phase”, and 3 terms concerning some interval, namely, “mid to late exponential phase”, “early to mid exponential phase” and “transition into stationary phase”. A contribution of the present work is that it not only includes terms depicting specific phases of growth but also collect terms related to intervals amongst two specific points, a practice commonly used by authors probably to get a better tracking of their own gene expression results. In this sense, terms such as “mid to late exponential phase”, “early to mid exponential phase” and “transition into stationary phase” were additionally incorporated in MCO.

Growth phase is a quality in the framework of BFO. Although BTO (BRENDA tissue/enzyme source) [35] describe growth phase cultures, we did not find terms to describe growth phases as such in OntoBee. Lag phase, acceleration phase (synonym of “transition into exponential phase”), exponential phase, retardation phase (synonym of “transition into stationary phase”), stationary phase, and phase of decline are clearly and unambiguously defined in terms of growth rates [36]. We added these terms as child nodes of growth phase. As we mentioned earlier, scientists frequently report experiments in a range of intermediate phases, but they do not report growth rates. This poses a complex ontological problem, because scientists do not precisely define these intermediate phases and it is possible that there is no way to unambiguously define them. It is very likely that a number of experiments reporting, for instance, “late exponential phase” indeed refer to different subintervals of exponential phase.

Thus, we roughly defined these intermediate phases in terms of the general ones. First, we defined “mid exponential phase” as a growth phase that is part of exponential phase and locates in the middle of the exponential phase. “early exponential phase” is a growth phase that is part of exponential phase and locates between acceleration phase and mid exponential phase. “late exponential phase” is a growth phase that is part of exponential phase and locates between mid exponential phase and retardation phase. “mid to late exponential phase” is a growth phase that comprises mid exponential phase and late exponential phase. “early to mid exponential phase” is a growth phase that comprises early exponential phase and mid exponential phase. “early stationary phase” is a growth phase that is part of stationary phase and locates right after retardation phase.

Despite their vague definitions, we consider that it is preferable to conserve all terms indicating growth subphases than to collapse all of them into their well-defined general term. This increases comparability of experiments. For instance, we believe that two experiments tagged with “late exponential phase” are more comparable than one that is tagged with “early exponential phase” and one with “late exponential phase”, even though the two experiments tagged with “late exponential phase” may have not been made in the same subinterval of exponential phase.

Based on this experience, and in order to increase reproducibility and comparability between experiments, we urge researchers to report more precise data related to the growth of cultures, similar to what has been done with the OD component, which, as described above, contain many specific values. As stated almost three decades ago by Neidhardt and colleagues: “*it should be noted that biochemical data are meaningful only if attention has been given to specify (1) the organism, (2) the growth environment, and (3) the state of growth*. *These parameters have a profound effect on biochemical results but often are not adequately documented in the reports of experiments”* [14].

### Inferences

Although the aim of recent curation was to find specific GCs terms, authors habitually use higher-level terms to indicate a specific condition. For instance, Lomovskaya *et al*. used “osmotic stress”, and “oxidative stress” terms to refer to the addition of NaCl 0.35 M or H_2_O_2_ 120 µM to the culture medium, respectively [37]. Regarding oxidative stress, we also observed that varying concentrations of H_2_O_2_, including 1, 100, 120, 500 µM, and 1 and 15 mM, as well as different compounds such as paraquat, are indistinctively associated with the same general term [25, 27, 37-45], revealing its amplitude and heterogeneity.

On the other hand, it is well known that, under certain growth conditions, the addition of chelating agents such as 2,2-dipyridyl or desferal to the culture medium, as well as deletion of the relA and/or spoT genes, reduce iron and ppGpp levels within the cell, respectively [26, 46-49]. Probably for this reason, most authors assume that a cell culture with those compounds or deletion of those genes will result, regardless of prevailing conditions, in iron or ppGpp depletion, thus they do not even experimentally measure the concentration of metabolites expected to vary or produce. Moreover, the decrease in iron or ppGpp is also related with more general terms like “low iron”, “iron depletion” and “stringent response”, respectively. Furthermore, these terms refer to some type of stress (see above).

A peculiar assumption, which underline the complexity of setting a unified vocabulary of growth conditions, is one that refers to trehalose as a stress condition [50] simply because, in *E. coli*, intracellular trehalose increases under high-osmolarity or starved culture conditions [51-54], while, in fact, trehalose does not harm but protects against these and other environmental stress such as desiccation, frost, and heat [55].

In brief, we found 24 terms inferring different kinds of stress or decompensation. However, we now only annotate either the specific compounds added to the medium or the genetic modifications explicitly indicated by the authors.

Our framework of annotation requires only material entities or measurable qualities that describe growth conditions. Since higher order concepts can be inferred from the real measurable entities that come into play in the laboratory, we included these higher order concepts in our ontology. We have future plans of making ontological relations that will allow us to do queries such as what are all of the agents of “oxidative stress” or what are all of the optical densities that indicate “high cell density”.

We believe that stress growth conditions do not refer strictly to a physiological state of the cell, but to a treatment applied to generate such physiological state. Therefore, although we found that oxidative stress was included in several ontologies (EO, MP, TO, EFO, OMIT, and NCIT) [56, 57] [30], we took the perspective of Plant Environment Ontology (EO). This ontology describes treatments that imply chemical stress. We took EO chemical stress treatment class along with its oxidative stress treatment and osmotic stress treatment child nodes. This does not possess a species-dependent compatibility problem, since the treatments are defined by their effects not by their agents nor by the species-specific responses they elicit. We extended this seed hierarchy to include the kinds of stress treatments used to study *E. coli*. We added nutrient availability stress treatment to describe nutrient limitation, nutrient depletion and nutrient excess treatments. We added as well the class temperature stress treatment to describe cold shock and heat shock. Desiccation and envelope stress were also included. It is worth noting that, in some cases, the definition of ontological relations between nutrient depletion stresses and its agents will be challenging, since we do not annotate the absence of any metabolite or property. In other cases, addition of a molecule implies the depletion of other, like in the case of chelating agents and metals.

We imported the NCIT class cell density along with its child nodes maximum cell density, mean cell density, and minimum cell density; and added high cell density and low cell density terms [https://github.com/NCI-Thesaurus/thesaurus-obo-edition]. We plan to relate these with specific optical densities. We imported oxygen content class from ZECO [**https://github.com/ybradford/zebrafish-experimental-conditions-ontology**] along with its child nodes hypoxia and hyperoxia. We merged this minimal hierarchy with aerobic environment and anaerobic environment terms from EO. We plan to relate these with specific concentrations of oxygen.

### GCs were defined by what they are, not by they are not

Similar to experiments that involved the addition of a particular supplement to the medium, experiments dedicated to analyze the influence of the absence of a certain compound in the genetic response are regularly performed. In these cases, authors generally use the prefix *absence of* or alternatively the addition of only a minus symbol at the end of each molecule name to explicitly indicate their absence, however, this would lead to long GCs descriptions. Moreover, the resulting phrases would surely contain attributes that were not considered in a given experiment, which in turn, may require to note the absence of *n* compounds that an experimental trial does not contain. Therefore, we would be defining a growth condition for something that is not. To avoid this, and similar to the decision of ignoring the large number of wild type genes of parent strains (see above), we decided not to annotate the absences. Hence, we obtained more concise phrases to indicate GCs composed only by the tangible features present in an experiment.

On the other hand, annotation of absences would lead to a counter-productive multiplicity of different terms, and consequently of different phrases referring to identical conditions. For example, regarding cells treated independently with distinct antibiotics [58], we observed that annotating the absence of each antibiotic in corresponding control samples lead to a proportional number of different phrases or overall growth conditions (Table 3, left and center columns), although all of them have the same meaning, and therefore may be expressed using only one phrase: “LB/ exponential phase”. Moreover, we would have greatly increased the number of terms that do not make biological sense, since annotating the absence of something equals nothing. Thus, by not indicating the absences, we describe only what an overall condition really is. Additionally, following this strategy, experimental variables are easily identified when comparing both control and experimental growth condition phrases. In this way, it can be distinguished when a particular experimental design involved the addition or remotion of either a gene, or compound (Table 3, right column), as well as the variation in a physical quality (see also display options in RegulonDB).

**Table 3.**
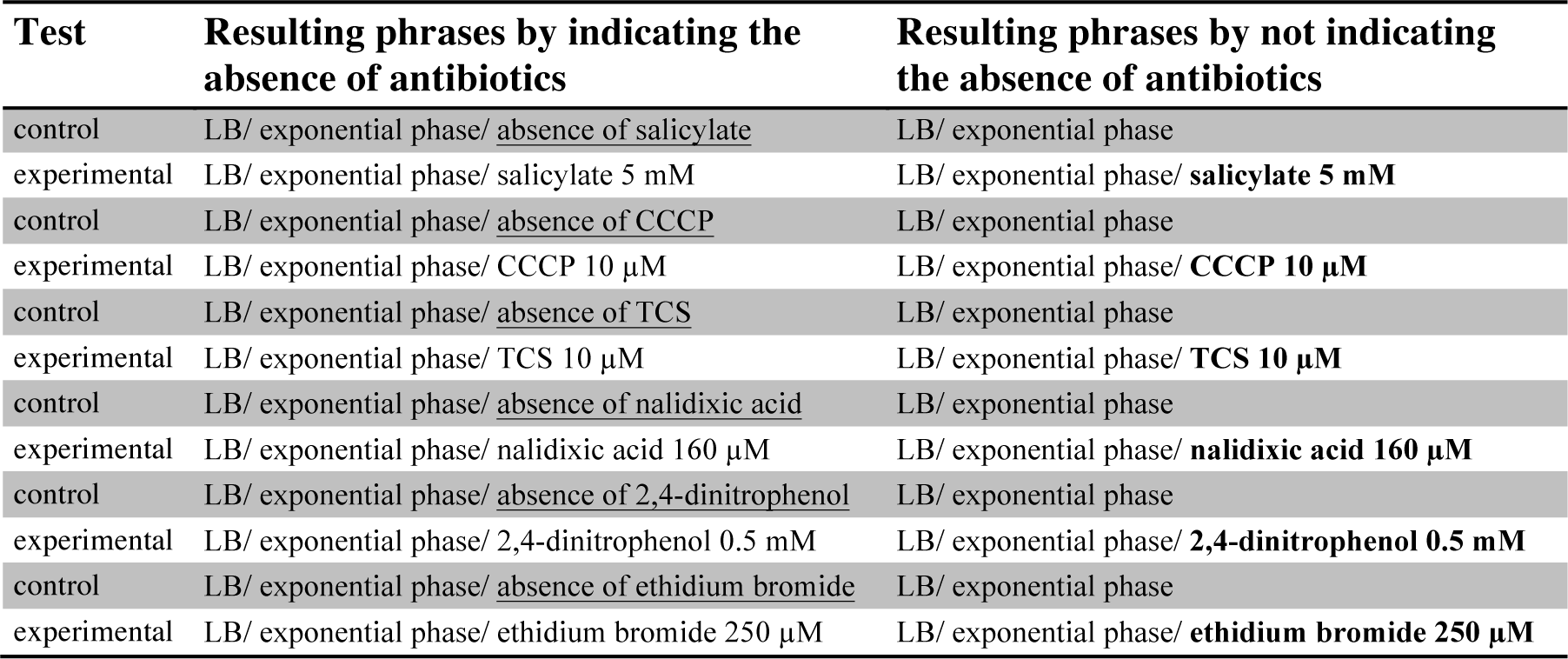
Comparison of growth conditions phrases that do or do not indicate the absence of a certain compound. The phrases were built from manual curation of an article that describe the genetic response of *E. coli* cells to the separate exposure to six different antibiotics [58]. Underlined terms specify the absence of tested antibiotics. Bold terms indicate the experimental variable for each control-experimental pair. CCCP, carbonyl cyanide m-chlorophenylhydrazone. TCS, tetrachlorosalicylanilide.

| Test | Resulting phrases by indicating the absence of antibiotics | Resulting phrases by not indicating the absence of antibiotics |
| --- | --- | --- |
| control | LB/ exponential phase/ <u>absence of salicylate</u> | LB/ exponential phase |
| experimental | LB/ exponential phase/ salicylate 5 mM | LB/ exponential phase/ **salicylate 5 mM** |
| control | LB/ exponential phase/ <u>absence of CCCP</u> | LB/ exponential phase |
| experimental | LB/ exponential phase/ CCCP 10 μM | LB/ exponential phase/ **CCCP 10 μM** |
| control | LB/ exponential phase/ <u>absence of TCS</u> | LB/ exponential phase |
| experimental | LB/ exponential phase/ TCS 10 μM | LB/ exponential phase/ **TCS 10 μM** |
| control | LB/ exponential phase/ <u>absence of nalidixic acid</u> | LB/ exponential phase |
| experimental | LB/ exponential phase/ nalidixic acid 160 μM | LB/ exponential phase/ **nalidixic acid 160 μM** |
| control | LB/ exponential phase/ <u>absence of 2,4-dinitrophenol</u> | LB/ exponential phase |
| experimental | LB/ exponential phase/ 2,4-dinitrophenol 0.5 mM | LB/ exponential phase/ **2,4-dinitrophenol 0.5 mM** |
| control | LB/ exponential phase/ <u>absence of ethidium bromide</u> | LB/ exponential phase |
| experimental | LB/ exponential phase/ ethidium bromide 250 μM | LB/ exponential phase/ **ethidium bromide 250 μM** |

Accordingly to curation details just described, and despite the drawbacks involved in notation standardization, which is necessary to construct the controlled vocabulary presented in this study, we collected 287 GCs terms. As stated above, these terms were joined to the terms recovered from the aforementioned datasets in RegulonDB, i.e. former GCs (64 terms), the effectors’ list (107 terms), and the TF notes (253 terms) to obtain a preliminary of 711 terms. However, this set yet included some repeated terms, that is to say, identical terms located in more than one dataset. Some of these terms were: "LB", "glucose", "paraquat", "37 ºC" and "exponential phase", each of which was initially present in at least 3 distinct datasets. Thus, after removing the repeated terms throughout all four datasets from RegulonDB, we were left with a total of 598 unique terms.

### Final constitution of the ontology of microbial growth conditions (MCO)

Similar to RegulonDB, Colombos, which also contains information on *E. coli* as well as other prokaryotic species, has also made an enormous effort to obtain the growth conditions terms of supported experiments. After analyzing the terms from both databases, we became aware that the repertoire of experimental conditions used, at least in *E. coli,* virtually remains the same regardless of the type of experiment, either using classic molecular biology or HT technologies. Therefore, GCs terms in RegulonDB are comparable to those in Colombos, thus building a unified ontology-based controlled vocabulary for these two databases is not only feasible, but will be highly rewarding given the large amount of classic experiments accumulated in RegulonDB and expression profiles in Colombos. To do this, we recovered 676 terms from Colombos that were not in RegulonDB. Accordingly, these terms were added to the unique terms we just obtained from RegulonDB leading to a unified vocabulary of 1274 distinct terms used to describe growth conditions used in experimental studies of *E.coli* K-12.

Grouped all the terms acquired from both databases, in addition to the terms recovered from other ontologies, constitute the first version of the Microbial Conditions Ontology (MCO) presented here. This ontology has 2765 classes, of which only the 21.5 % are original MCO classes, 75 % are classes from ChEBI, and 3.5 % come from other supporting ontologies.

### The ontology and GCs in RegulonDB

Having standardized and defined the set of elementary terms in the ontology, we built the composed growth condition annotation phrases, of the specific curated experiments, in RegulonDB. These phrases are as follows:

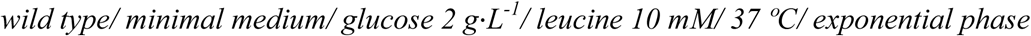

As mentioned before, these composed phrases describe material entities and measurable qualities of the medium in which cells were grown, as well as the identity of the cell type. We can only have access to what authors report in the literature (Fig. 2), therefore, sometimes we cannot fill all slots of required items of information defined in our annotation framework. Despite this, the order of reported elements will be constant in phrases (see Methods section).

For each experiment, only one GC phrase will be used to annotate either the control- or the experimental growth conditions. As described in Table 3, the experimental variable becomes evident when comparing two or more phrases containing common terms. In the example shown below, the variable is leucine as it is the element that is only part of the description of the experimental condition. Regardless their role in experimental design, either basal or variable, all terms are included in the ontology. In fact, it is only throughout the queries at regulondb.ccg.unam.mx, when it can be seen the precise role that a particular term is playing in any of the displayed growth conditions.

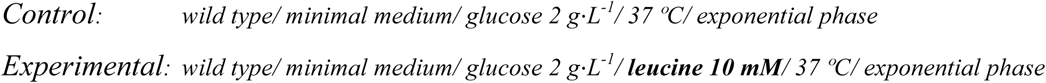

The elementary terms for growth conditions in MCO have been integrated into RegulonDB and linked to the GCs composed terms, which in turn are linked to their affected objects as genes, transcription factors, etc. The ontology can be accessed in the “integrated views and tool” menu. Furthermore, the user can navigate through the terms list in the ontology browser, and when a term is selected, the GC phrases are displayed in the web page with the information about the experiments and their affected objects. The user can do a search using the search text box for a very specific term as “glucose 1 mM”, or less specific terms, such as “glucose”, or “carbohydrates”. A list of composed terms (phrases) representing the GCs containing the searched term will be shown as the first result. If one of the obtained GC is chosen through a click, a web page containing all the genes affected by the GC is shown. In that page, details of each record such as the effect on each gene caused by the GC, the TF and the promoter affected by the GC will be shown. Also the evidence, method and references supporting the effect of the GC on gene expression are displayed. The searches can be directed towards the set of GCs used as variables in experiments or more generally towards all GCs. To do that, a button is displayed in the search section that indicates to do a search of a specific term used as variable (term as variable) and a button to indicate that the search is done both in the variables and in the basal properties terms of the GC (all terms).

On the other hand, in the pages of gene, TU and TF there is a new link to GCs affecting such elements. The links points to a web page listing all GCs affecting the object in addition to specific details related to each GC.

## Discussion and conclusions

Once we had the collection of elementary GCs terms, we started to build phrases describing the GCs in which each experiment was done. We linked a pair of phrases describing two experiments, the test and control, to the genes whose expression was affected. When we started years ago to annotate GCs in RegulonDB, we only annotated the terms related to the molecule, physical quality or growth phase that was the contrasting variable analyzed in the experiments; as a consequence, in those cases we do not have a complete phrase describing the GCs. Such data will remain incomplete in our database until the annotation of these elements is re-annotated in the future.

On the other hand, we have recently initiated assisted curation processes, that consist in the curation of selected phrases or paragraphs that a semiautomatic system, developed in our research group, extracts from complete articles for the curator to proceed [59]. One strategy implemented to find informative phrases is to identify the sentences that contain a relationship between the GCs, the target gene and the effect that is caused by the GCs. Using this strategy, we are able to identify only the terms that represent the compounds that are precisely variable in the experiments. Therefore, the data so far obtained in this way is currently also incomplete. We will need to reevaluate what strategy to follow in our continued assisted curation to identify all the elements of GCs defined in MCO.

The scope of the ontology is clear and succinctly worded: an ontology of growth conditions of microbial organisms. However, kinds of entities that compose growth conditions are very diverse. According to BFO, in our developed framework of annotation, growth conditions are composed of qualities, material entities, and processes. As a result of this diversity of entities, our ontology turned out to be composed of imported terms from other ontologies, but sorted differently. Thus, only a fraction of our ontology is composed by collected terms from both RegulonDB and Colombos. However, since our ontology includes the strain and genetic background of both the experiment and control, it goes beyond strictly growth conditions, to encompass the set of minimal properties necessary and sufficient to support the reproducibility and comparison of the experimental setting of microbial research on gene regulation. These properties are certainly fundamental for their description, but as mentioned, insufficient to guarantee reproducibility given the lack of growth rate, and in a broader sense, the lack of all additional experiments performed to link these conditions to the consequences in gene expression and the discovery of regulatory mechanisms.

Despite the fact that most of our terms come from other ontologies, we can argue that our ontology is orthogonal to other ontologies invoking the methodological principle proposed in [5] of adequatism. This principle accepts the need of alternative views of reality. It was originally proposed to allow the construction of theories that reflect two dimensions of plurality: the opposition of different levels of granularity and the opposition between objects and processes. Here, however, we make a slice through these kinds of oppositions and use entities of different levels of granularity, as well as entities of objects and processes. The plurality in perspective is embedded in the general idea that, besides being a quality, an object, a role, a process, or a spatiotemporal region, these entities also define experimental conditions. As we used the post-composition approach [15], the relationship between the entities and conditions of or our ontology is realized in the annotation process. Due to the multi-dimensional nature of the growth conditions description, the potential number of different conditions can be astronomical. Therefore, we used the post-composition approach in order to keep simplicity without compromising the comprehensiveness of the annotation.

Growth condition terms described in this study come from a small sample of the total of papers currently supporting RegulonDB, so that we are aware that as this work continues new terms will be surely added, but the logic and structure here defined should be in any case, minimally modified. In fact, other than the adequacy of the specific terms in our ontology, and some enhancements in cell types (differentiation such as sporulation) and anatomy of the population structure (i.e. biofilms), this framework can also be used to support a better description of the knowledge on gene regulation in all the microbial world. This comprehensiveness is better appreciated when compared to the minimalist approach of describing only the -variable/absence of variable-contrasting description, used for instance in Colombos. First of all, the comprehensiveness we propose can, in principle, enable new -*in silico* built-experiments by performing novel comparisons between pairs of conditions that have not been yet compared. Second, as mentioned earlier, the search for comprehensiveness derives from the ontological requisites of definitions as the set of minimal and sufficient conditions for a given definition. And third, this brings up to our attention the lack of explicitly stating growth rate in the literature we curate, in spite of its relevance as mentioned by Neidhardt decades ago, and more recently by Hwa in theoretical developments of microbial physiology [60].

The applications of the ontology here described, together with its unified vocabulary, is an essential part in the foundation for the comparison and integration of the large amounts of knowledge on gene regulation coming from different sources, and methods, particularly classic molecular biology as well as high throughput methods, a current effort in our laboratory. For instance, this ontology is used in our current curation of HT-binding experiments with ChIP-exo, ChIP-seq, or gSELEX permitting us to assign the function of a site when a change in expression is found in Colombos; it is also the basis to strengthen the confidence level of a given interaction, binding site, or any other piece of knowledge, when supported by different methods performed under the same conditions [61].

As growth conditions are more exhaustively curated, we will be able to gain knowledge from the currently genotypic-centered *E.coli* transcriptional regulatory network (with all interactions without knowing when they are active, with some clearly never co-occurring), to its mapping into the phenotypic networks active under particular conditions.

As microbial knowledge at the genomic level will proceed in the future of research, there is no doubt that an ontology of growth conditions and the experimental setting of changes of gene expression, together with an equally comprehensive ontology of the machinery of gene regulation, are two essential complementary pieces to provide a solid foundation of past, current and future microbial physiology.

## Funding

This work was supported by the National Institutes of Health [grant number R01GM110597-3] and FOINS CONACyT Fronteras de la Ciencia [project 15]. CMA is a doctoral student from Programa de Doctorado en Ciencias Biomédicas, Universidad Nacional Autónoma de México (UNAM) and received fellowship 576333 from CONACYT.

## Acknowledgements

Authors wish to thank Alberto Santos-Zavaleta, Cecilia Ishida-Gutiérrez, David A. Velázquez-Ramírez and Sara B. Martínez-Luna for meaningful discussion during curation process. We also thank Víctor M. Del Moral-Chávez and César Bonavides-Martínez for their technical support.

